# Glyoxal as alternative fixative for single cell RNA sequencing

**DOI:** 10.1101/2021.06.06.447272

**Authors:** Josephine Bageritz, Niklas Krausse, Schayan Yousefian, Svenja Leible, Erica Valentini, Michael Boutros

**Affiliations:** German Cancer Research Center (DKFZ), Division Signaling and Functional Genomics and Heidelberg University, Department of Cell and Molecular Biology, Medical Faculty Mannheim and BioQuant, Im Neuenheimer Feld 580, 69120 Heidelberg, Germany; Heidelberg University, Centre for Organismal Studies, Research group Stem Cell Niche Heterogeneity, Im Neuenheimer Feld 230, 69120 Heidelberg, Germany

**Keywords:** single cell RNA sequencing, transcriptomics, cell fixation, glyoxal

## Abstract

Single cell RNA sequencing (scRNA-seq) has become an important method to identify cell types, delineate the trajectories of cell differentiation in whole organisms and understand the heterogeneity in cellular responses. Nevertheless, sample collection and processing remain a severe bottleneck for scRNA-seq experiments. Cell isolation protocols often lead to significant changes in the transcriptomes of cells, requiring novel methods to preserve cell states. Here, we developed and benchmarked protocols using glyoxal as a fixative for scRNA-seq application. Using Drop-seq methodology, we detected high numbers of transcripts and genes from glyoxal-fixed *Drosophila* cells after scRNA-seq. The effective glyoxal fixation of transcriptomes in *Drosophila* and human cells was further supported by a high correlation of gene expression data between glyoxal-fixed and unfixed samples. Accordingly, we also found highly expressed genes overlapping to a large extent between experimental conditions. These results indicated that our fixation protocol did not induce considerable changes in gene expression and conserved the transcriptome for subsequent single cell isolation procedures. In conclusion, we present glyoxal as a suitable fixative for *Drosophila* cells and potentially cells of other species that allows high-quality scRNA-seq applications.

## Introduction

The development of single cell RNA sequencing (scRNA-seq) methods has opened new analytical avenues in the molecular life sciences (Picelli, 2017; Aldridge and Teichmann, 2020). Genome-wide transcriptomic data is now routinely generated with single cell resolution on plate-(Picelli et al., 2013) or microfluidics-based systems (Macosko et al., 2015; Klein et al., 2015). Despite a rapidly expanding number of improved technologies, the overall workflow remains similar. Following separation, single cells are lysed in individual reaction chambers. The released mRNA is converted into cDNA libraries, PCR-amplified and further processed for high-throughput sequencing. Most enhancements of scRNA-seq technologies focus either on different aspects of the sequencing library preparation or downstream analysis tools. However, also single cell sample preparation is of central importance. Whether working with cell lines or primary tissue, optimizing and tailoring the cell isolation protocol for the specific sample type is important for the overall single cell data quality. The aim is hereby to fully disaggregate the sample into single cells without compromising their viability or integrity, which minimizes the level of free RNA from dead, dying or damaged cells and thus reducing experimental noise. While single cell RNA sequencing experiments should be carried out without much delay in order to prevent degradation of RNA and thus changes in the transcriptome, in reality, the experimental design often requires flexibility in the sample collection process. Single nuclei RNA sequencing (nucSeq) from frozen tissue samples offers a suitable way to bypass this constraint. However, the lower transcript counts obtained from nucSeq (Habib et al., 2017; Bakken et al., 2018) can constitute a drawback of this method, especially when working with model organisms with low RNA content, such as *Drosophila*.

Sample fixation has the potential to preserve the transcriptome and ease the scRNA-seq experiment at the same time. However, despite the strongly increasing number of scRNA-seq studies (Svensson et al., 2018), protocols with fixed single cells are only used to a limited extent (Karaiskos et al., 2017; Alles et al., 2017; Attar et al., 2018; Thomsen et al., 2015; Wang et al., 2020; Wohnhaas et al., 2019; Denisenko et al., 2020; Van Phan et al., 2020). Among the observed disadvantages are a reduced library complexity with lower number of detected transcripts (Alles et al., 2017; Attar et al., 2018; Van Phan et al., 2020) and higher level of ambient RNA (Wohnhaas et al., 2019). A comparative analysis of alternative fixation methods, however, has not yet been performed.

Here, we performed glyoxal fixation on *Drosophila* and human cell lines and analyzed their transcriptome using Drop-seq. The gene expression profiles of unfixed and glyoxal-fixed samples showed an overall high similarity for both species. In particular, glyoxal fixation of *Drosophila* cells resulted in high quality scRNA-seq data with a low fraction of mitochondrial encoded RNA, and high number of detected genes and transcripts. Our glyoxal fixation protocol has the potential to increase both quality and comparability of scRNAseq data in particular for *Drosophila* samples and other model organisms with low RNA amount.

## Methods

### Cell line fixation for scRNA-seq

Kc167 cells were cultured in Schneider’s media supplemented with 10% heat-inactivated FCS at 25°C in T75 cell culture flasks with solid caps. HEK 293T cells were cultured in DMEM supplemented with 10% fetal calf serum (FCS) and 1% penicillin/streptomycin at 37°C and 5% CO_2_ in T75 cell culture flasks with filter caps. For preparing a single cell solution, Kc167 cells were mechanically detached, transferred to a 15 ml tube and centrifuged at 260 rcf for 5 min at room temperature. The cell pellet was washed once with filtered PBS, resuspended in filtered PBS and filtered through a 20 µm cell strainer. HEK 293T cells were washed with filtered PBS (0.22 µm filter) and enzymatically detached with 2 ml 1x TrypLE. The reaction was inactivated with 10 ml filtered PBS. Detached cells were filtered through a pre-equilibrated 40 µm cell strainer. Cell concentrations were determined with a disposable Neubauer hemocytometer (C-Chip DHC-N01). Unfixed cells have been directly processed by scRNA-seq. For glyoxal fixation, 5 × 10^6^ cells per condition and cell line were resuspended in 1 ml 3% glyoxal mix pH 4 (Richter et al., 2018) and incubated for 1 h on ice. Next, the fixed cell solution was centrifuged at 260 rcf for 5 min at 4°C, washed twice with filtered PBS and filtered through a pre-equilibrated 40 µm (HEK 293T) or 20 µm (Kc167) cell strainer. HEK 293T and Kc167 single cells were equally mixed to a final cell concentration of 100 cells/µl and used for scRNA-seq.

### Single cell RNA sequencing by Drop-seq

Single cell transcriptomic data was generated with Drop-seq following available protocols (Macosko et al., 2015; Bageritz and Raddi, 2019). In brief, cells and barcoded beads (ChemeGene) were co-flowed into a T-junction polydimethylsiloxane (PDMS) microfluidic device (FlowJem) and co-encapsulated in 125 μM droplets. High quality emulsions were broken by perfluorooctanol and reverse transcription of captured polyadenylated mRNA was performed. Subsequently, barcoded beads were incubated with Exonuclease I to remove excess primers, and cDNA was amplified with 14 PCR cycles from 2x 2,000 beads (replicate 1) and 4,000 beads (replicate 2), respectively. A 0.6 ratio of AMPure beads (Agencourt) was used to purify cDNA libraries, which were eluted in 10 μl water. Final libraries were prepared using the Illumina Nextera XT kit. Paired-end sequencing was carried out with the Illumina HiSeq2500 instruments at the DKFZ Genomics and Proteomics Core Facility (Heidelberg, Germany). Experiments have been performed in two biological replicates. For the first biological replicate, technical replicates were generated during PCR amplification, while the second biological replicate included technical replicates from the unfixed sample from two different Drop-seq runs. Individual libraries were prepared for each replicate and sequenced. For downstream analysis individual technical replicates were merged and analyzed together.

### Processing and quality assessment of scRNA-seq data

Sequencing data was processed as described by Macosko and colleagues (Macosko et al., 2015). An interface to the used R functions was implemented in our in-house Galaxy^1^ server following the default settings described in detail in the Drop-seq computational cookbook v.1.2 accessed at http://mccarrolllab.org/dropseq/ and described in detail (Macosko et al., 2015). The reads were aligned to the *Drosophila* reference genome (BDGP6 v.91 (GCA 000001215.4) and GRCh37.87 (hg19))) using STAR v.2.5.2b-0 with the default parameters and showed at least 70% uniquely mapped reads for all samples. The CollectRnaSeqMetrics tool from Picard v2.18.2 (Broad Institute 2018). was used to collect metrics about the alignments. The cell number was estimated by plotting the cumulative fraction of reads per cell against the sorted cell barcodes (decreasing number of reads) and determining the point of inflection. The generated digital gene expression matrices were then further analyzed using the R package Seurat v3.1.5 (Satija et al., 2015). We kept cells with a minimum of 200 detected genes, analyzed the fraction of mitochondrial encoded RNA, and removed outlier cells from further analysis. The chosen threshold was <5% for the *Drosophila* samples and <10% for the unfixed and <20% for the glyoxal-fixed human sample. The level of species-mixed cell barcodes was assessed, and cells with < 100% purity assigned as mixed population and discarded from further analysis.

### Data normalization

As samples have been sequenced on 5 flow cell lanes, they showed batch effects mainly originating from differences in sequencing depths. We therefore decided to normalize them with the following steps: From each BAM file we first extracted the reads belonging to the real cells as determined from the knee plot, we then randomly sampled a fraction of reads from each extracted BAM file to have the same average number of reads per cell among the different samples. The library with the lowest average number of reads per species was taken as reference. *Drosophila* samples were normalized to 12,697 reads per cell and human samples to 24,791 reads per cell. For both operations, we used pysam, a python wrapper around the samtools package (https://github.com/pysam-developers/pysam, Li et al., 2009).

### Evaluation of library complexity

The sequencing saturation level is a measure of the fraction of library complexity that was sequenced in a given experiment and has been calculated for the normalized libraries using the following formula:

Sequencing saturation = 1 - (n_deduped_reads / n_reads), where n_deduped_reads is the number of transcripts (UMI) and n_reads is the number of reads.

### Sample comparison

To compare the biological replicates and the cells treated with glyoxal and untreated for both human and *Drosophila*, we computed the gene expression correlation of the genes in common. The gene expression was normalized by calculating the average UMI expression for each gene and converting it to average transcript per million (ATPM). We then calculated the Pearson’s correlation coefficient (R) based on the normalized expression values. Single cell expression data was aggregated to ‘pseudobulk’ data and the 100 most expressed genes were compared between unfixed and glyoxal-fixed samples.

## Data Availability

The generated scRNA-seq count matrices are deposited in Gene Expression Omnibus and accessible through accession number GSE163736.

## Code availability

Single cell transcriptome analyses have been performed with R 3.6.3 (using R studio). The code to reproduce the figures including the normalization steps will be available on request.

## Results

### Effects of glyoxal fixation on scRNA-seq performance

Glyoxal is a dialdehyde used as fixative for immunohistochemistry for many years (Dapson, 2007) and was recently shown to present a promising alternative to the commonly used paraformaldehyde (Richter et al., 2018; Channathodiyil and Houseley, 2021). Compared to PFA glyoxal has a lower toxicity, a faster fixation speed and shows little tendency to crosslink under certain pH conditions. Those features also make glyoxal a promising fixative for single cell transcriptome studies. RNA molecules would be quickly preserved without molecule leakage and impaired purity. It would also allow high amounts of RNA recovery without the need to reverse crosslink the samples. To evaluate glyoxal as fixative for scRNA-seq applications we performed head-to-head comparison of unfixed (blue) and glyoxal-fixed (green) *Drosophila* (Kc167) and human (HEK 293T) cells. Glyoxal fixation was performed by buffering the solution to pH 4, a condition that inhibits crosslinking (Dapson, 2007). As described before (Macosko et al., 2015), mixed species populations were subjected to scRNA-seq and the obtained transcriptomic data was thoroughly compared (Fig.1 a).

**Figure 1.**
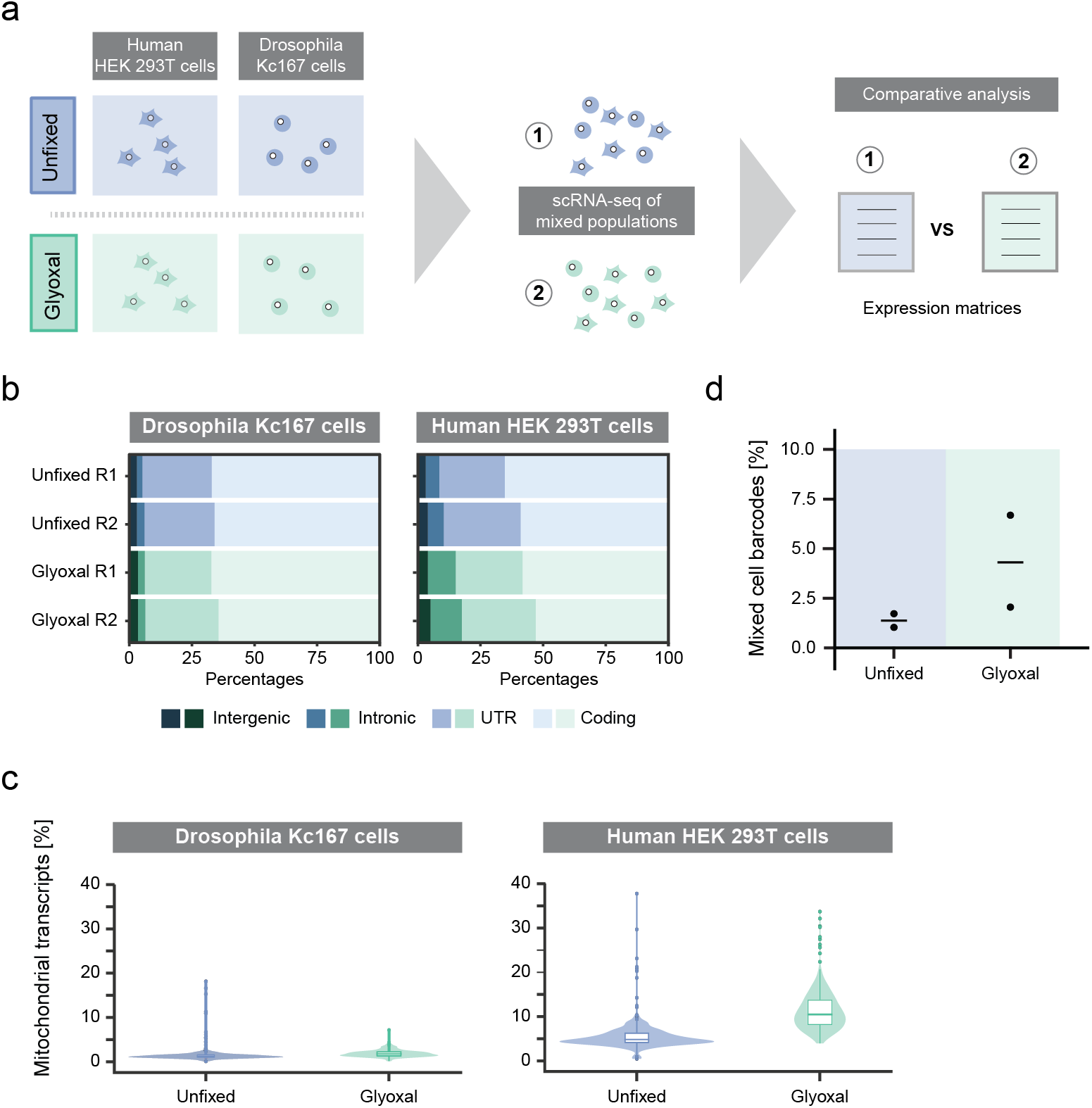
Effect of glyoxal fixation on cell integrity, RNA quality and purity. (**a**) Workflow to assess quality of glyoxal-fixed human and Drosophila cell lines for single cell RNA transcriptome studies. Single cell suspension from different species were mixed equally, subjected to Drop-Seq and their transcriptome compared by downstream analysis. (**b**) Proportion of reads mapped to coding, UTR (untranslated region), intronic, and intergenic regions. Two biological replicates were analyzed per condition for each cell line. (**c**) Fraction of mitochondrial transcripts out of total mRNA transcripts per condition and cell line. (**d**) Percentage of mixed cell barcodes detected per data set. Horizontal line indicates the mean.

In order to preserve the transcriptome of a cell, a fixative has to act fast to prevent loss of cytoplasmic mRNA due to leakage of RNA molecules through pores in the cell membrane. In case of RNA leakage, the relative abundance of intron-containing nascent transcripts of the nuclear compartment would increase due to its more protected location. By analyzing the distribution of the transcript coverage in the *Drosophila* data, we found the average read mapping to distinct transcript regions highly comparable between glyoxal-fixed and unfixed samples (Fig. 1b, left). In contrast, glyoxal fixation of human cells introduced minor changes in the transcript coverage. Here, we found a slight increase in the average read number mapping to intronic regions from 6 % to 12 %, while the average fraction of mature transcripts (UTR, coding) was reduced from 91 % to 84 % upon glyoxal fixation (Fig. 1b, right). Similarly, determining the fraction of mitochondrial RNA within each cell can be used as an indicator for loss of cytoplasmic RNA in cells (Ilicic et al., 2016). In *Drosophila* cells, the proportion of mitochondrial RNA detected in unfixed and glyoxal-fixed samples was again comparable (Fig. 1c, left). A median of less than 2 % mapped mitochondrial transcripts has been found in both, unfixed and glyoxal-fixed cells. In contrast, glyoxal fixation of human cells led to an increase in mapped mitochondrial transcripts with a median fraction of 4.8 % in unfixed and 10.5 % in glyoxal-fixed cells (Fig. 1c, right). Next, we assessed the single cell purity of the samples and obtained a similar ratio of mixed transcriptomes between unfixed and glyoxal-fixed cells (Fig. 1 d). Low level of ambient RNA is apparent in the overlaid kneeplots (Additional File 1, Supp. Fig. 1a), showing a comparable fraction of reads in unfixed and glyoxal-fixed cells, while empty droplets only captured little amount of ambient RNA.

Taken together, we show that glyoxal fixation is compatible with downstream single cell RNA sequencing methodology and that RNA molecules seem to be better fixed by glyoxal in *Drosophila* cells than they are in human cells.

### Preservation of cytoplasmic RNA molecules upon glyoxal fixation

Next, we examined the impact of glyoxal fixation on cytoplasmic mRNA transcripts. We observed a shorter average cDNA size for the species-mixed glyoxal libraries compared to the unfixed samples (Additional file 1, Supp. Fig. 1b), which indicates RNA degradation and/or fragmentation in Drosophila and/or human cells. To further assess the effect of RNA leakage and degradation/fragmentation, we analyzed the level of detected genes and transcripts. For this purpose, we normalized the data for differences in sequencing depth (Additional file 2, Supp. Fig. 2a) and found the samples to have a mean library saturation of about 50% – 70% (Additional file 2, Supp. Fig. 2b). Notably, the unfixed and glyoxal-fixed Drosophila samples showed a similar mean saturation level, while the human glyoxal sample showed an almost 20% higher mean saturation than the unfixed human samples indicating lower library complexity. Regardless of the species or experimental condition the biological replicates correlate well (R ≥ 0.88 for Drosophila and human data, Additional file 2, Supp. Fig 2c) allowing us to thoroughly compare the experimental conditions.

In *Drosophila*, the detected gene and transcript counts show a similar distribution in fixed and unfixed cells (Fig. 2a). The median recovery of genes and transcripts for unfixed (1,398 genes/cell and 4,577 transcripts/cell) and glyoxal-fixed (1,408 genes/cell and 4,000 transcripts/cell) are highly comparable. In contrast, the human sample showed a lower variation in detected gene and transcript expression and a reduction in the median number of genes and transcripts upon glyoxal fixation (Fig. 2b). In unfixed human cells, we detected a median of 3,426 genes/cell and 11,142 transcripts/cell, whereas in glyoxal-fixed cells that number was reduced to 2,684 genes/cell and 6,812 transcripts/cell, respectively.

**Figure 2.**
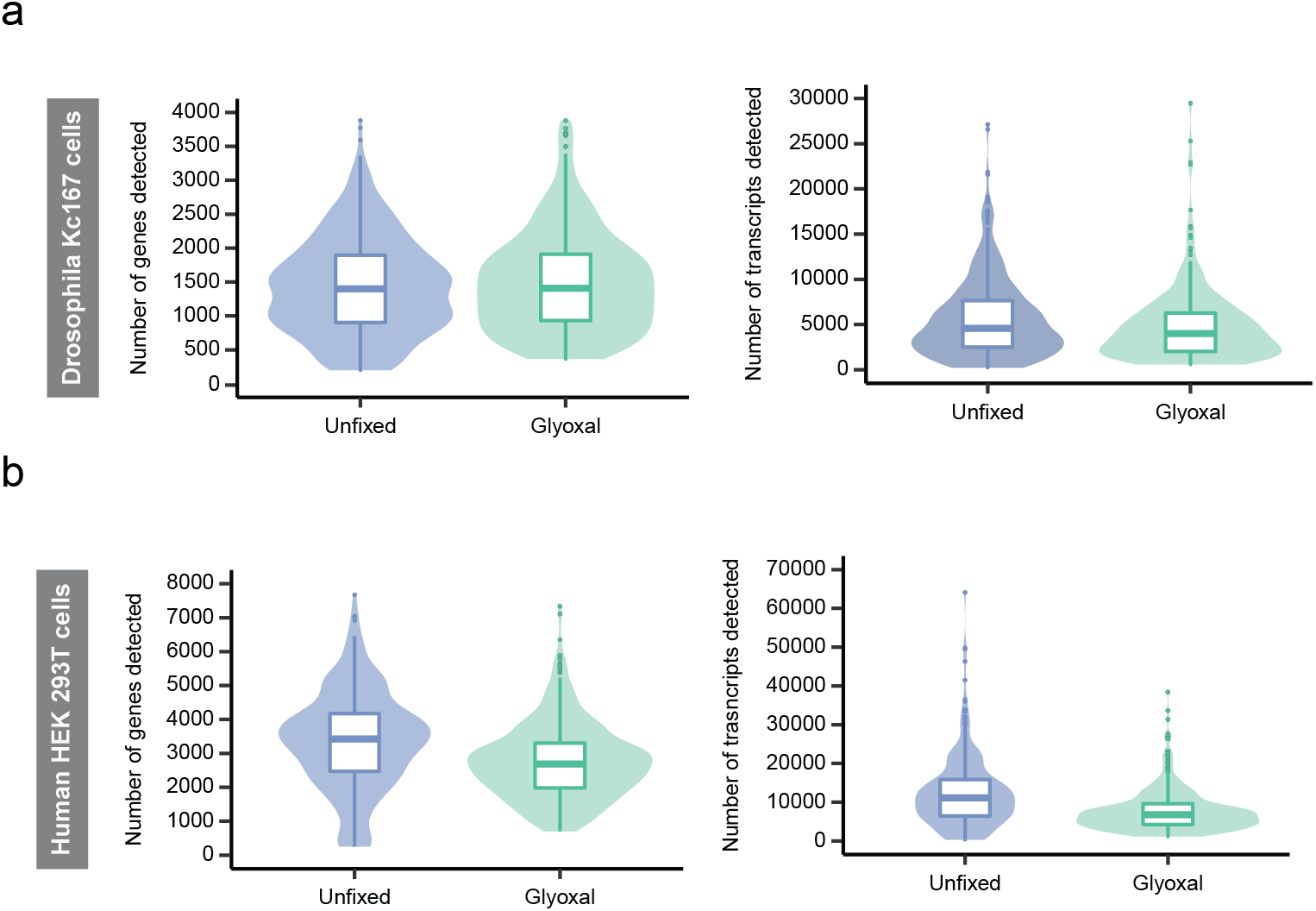
Library complexity in unfixed and glyoxal-fixed samples. (**a-b**) Number of detected genes (left) and transcripts (right) for unfixed and glyoxal-fixed single cell solutions obtained from *Drosophila* (**a**) or human (**b**) cell lines. Two biological replicates were analyzed per condition and cell line. Box plots show median and 25th and 75th percentiles.

In summary, our data suggest that glyoxal effectively preserves RNA molecules of *Drosophila* cells, while human cells seem to be more prone to cytoplasmic RNA leakage and RNA degradation and/or fragmentation resulting in lower number of detected genes and transcripts.

### Glyoxal fixation in *Drosophila* and human cells preserves the mRNA

In order to examine the effect of glyoxal fixation on the entire transcriptome, we aggregated single cells to ‘pseudobulk’ data and compared their gene expression profiles (Fig. 3a). By analyzing the set of commonly expressed genes (8,040 genes in *Drosophila* cells, 17,092 genes in human cells), we showed a high correlation between unfixed and glyoxal-fixed samples (R = 0.95 for the *Drosophila* data, R = 0.94 for the human data). This allows the conclusion that the transcriptomes of unfixed and glyoxal-fixed *Drosophila* and human cells are highly similar when analyzed as ‘pseudobulk’ data sets. We then extracted the top 100 highly expressed genes from both conditions and determined their overlap (Fig. 3b). Comparing the unfixed and glyoxal-fixed samples showed with 95 genes in *Drosophila* and 90 genes in human a high overlap, which is similarly seen by comparing the respective technical replicates (Additional file 3, Supp. Fig. 3) indicating that glyoxal fixation does not change gene expression profiles substantially.

**Figure 3.**
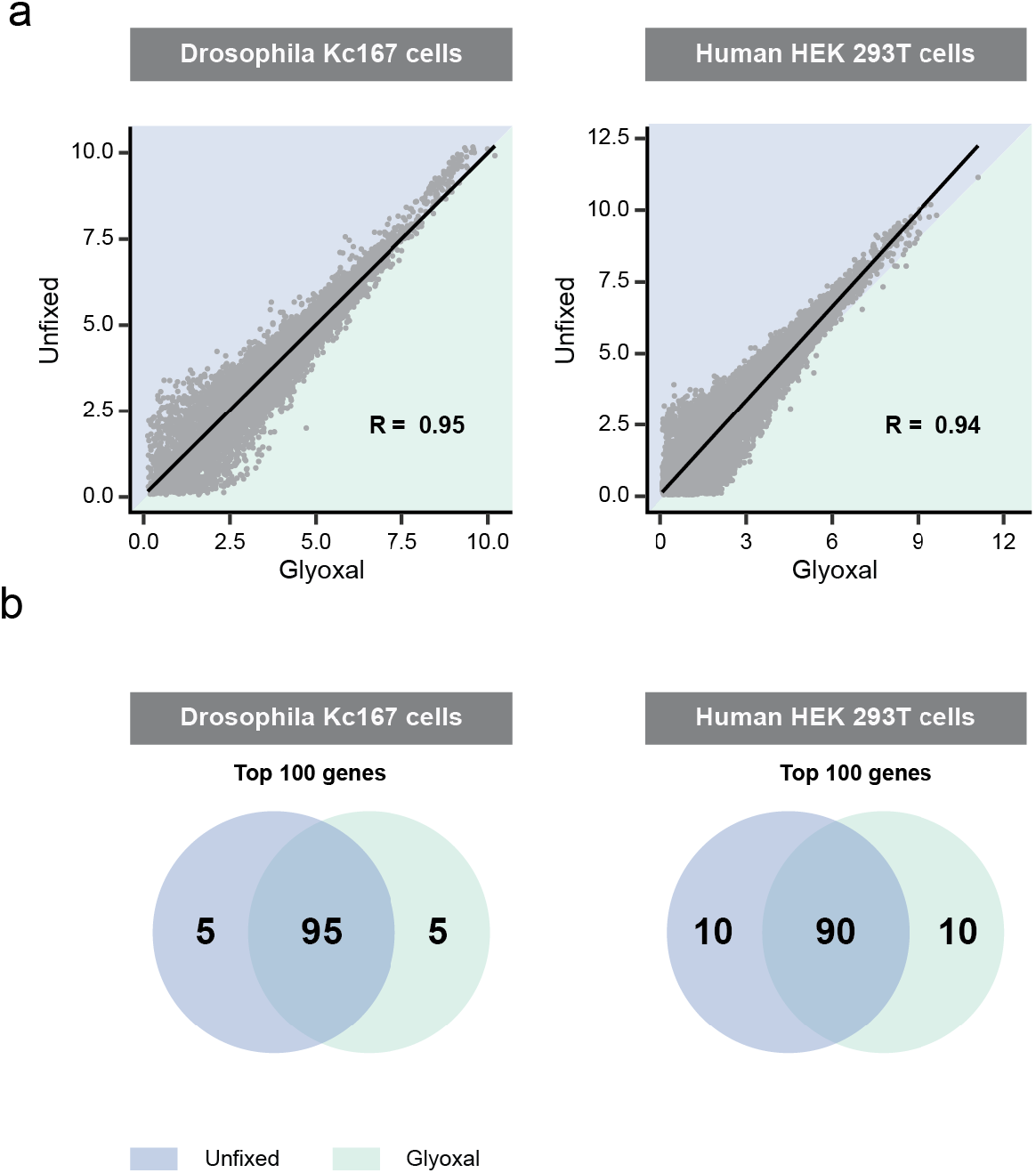
Transcriptome similarities of unfixed and glyoxal-fixed *Drosophila* and human cells. **(a)** High average gene expression correlation between unfixed and glyoxal-fixed libraries. Average gene expression correlation of shared genes between pooled unfixed and glyoxal-fixed Drosophila and human cell lines. Data is shown as normalized ATPM. R indicates Pearson correlation coefficient. **(b)** Overlap between top 100 highly expressed genes for unfixed and glyoxal-fixed *Drosophila* (left) and human (right) cell lines depicted in Venn diagrams.

Taken together, in this report we have identified glyoxal as a suitable fixative for scRNA-seq applications. We found glyoxal fixation to be compatible with high-throughput single cell sequencing with fixed samples maintaining a preserved transcriptome of high purity, and library complexity.

## Discussion

Glyoxal has successfully been used as fixative for detecting single mRNA molecules by in-situ stainings and also the applicability to perform bulk RNA-sequencing has been shown (Richter et al., 2018; Channathodiyil and Houseley, 2021). Here we report the additional use of glyoxal as fixative for scRNA-seq. Although, previous studies have presented different fixatives for scRNA-seq applications (Karaiskos et al., 2017b; Alles et al., 2017; Attar et al., 2018, Van Pahn et al., 2020, Thomsen et al. 2016), we have found glyoxal fixation to be highly advantageous in many key aspects when it comes to the preservation of transcriptomic data.

Generally, in scRNA-seq experiments only a small fraction of the transcriptome is captured and so downstream analysis can be quite challenging. High number of detected genes and transcripts is hence aimed for, especially in tissues or model organism with lower number of detected mRNA transcripts in scRNA-seq experiments such as *Drosophila* (Karaiskos et al., 2017; Bageritz et al., 2019). In contrast to other fixatives, glyoxal fixation did not compromise the level of detected transcripts and genes in *Drosophila* Kc167 cells. Library complexity in human HEK 293T cells was reduced upon glyoxal fixation, but not to an extent seen for nucSeq experiments (Habib et al., 2017) and to a similar level than methanol fixation (Alles et al., 2017; Chen et al., 2018). Differences in fixation speed impacting RNA leakage from the cytoplasm and RNA degradation most likely explain the observed quantity loss in human cells. Titrating the pH of the glyoxal mix as previously done for optimizing the immunostaining protocols (Richter et al., 2018) might overcome this limitation for scRNA-seq experiments. Importantly, glyoxal fixation in both human and *Drosophila* cells yields transcriptome data similar to that from unfixed cells with technical noise greater than any effect of glyoxal. This also indicates that glyoxal fixation did not compromise transcript accessibility for library preparation and sequencing.

Isolating single cells from solid tissue can be challenging as dissociation protocols need to be optimized to avoid cell damage and the resulting increased levels of free RNA while at the same time facilitating unbiased recovery of single cells in sufficient quantity. Fixation of the tissue prior to single cell dissociation could solve this problem. However, there is currently no fixative available that can be efficiently used for tissue fixation prior to the application of any lengthy dissociating protocols. In this regard, it will be interesting to analyze the single cell recovery of glyoxal-fixed primary tissue. Glyoxal might additionally be useful to uncouple sample collection from further processing by allowing sample storage. This can be useful for clinical samples or in experiments that require a large amount of starting material. Overall, the data presented here highlights glyoxal as a suitable fixative for scRNA-seq experiments.

Improved sample preservation will aid in the generation of high-quality data sets for scRNA-seq. Here we present with glyoxal a comparably cheap, easy to handle and most of all effective fixative, which can be easily implemented in existing experimental work flows.

## Acknowledgements

We thank Bianca Balzasch for technical assistance with the sample preparations. We thank the High Throughput Sequencing Unit of the DKFZ Genomics and Proteomics Core Facility for providing excellent next-generation sequencing services. We thank C. Previti for implementing the DropSeq computational cookbook on the Galaxy Server. We thank our colleagues Florian Heigwer and Jan Gleixner for proof-reading the manuscripts. J.B. was supported by a research stipend from the Fritz Thyssen Foundation. This project was supported by funds from the DFG (SFB1324) and the European Commision (ERC Synergy Project DECODE).

## Notes

### Competing Interest Statement

The authors have declared no competing interest.

