## Supplementary figures and figure legends for "Glyoxal as alternative fixative for single cell RNA sequencing"

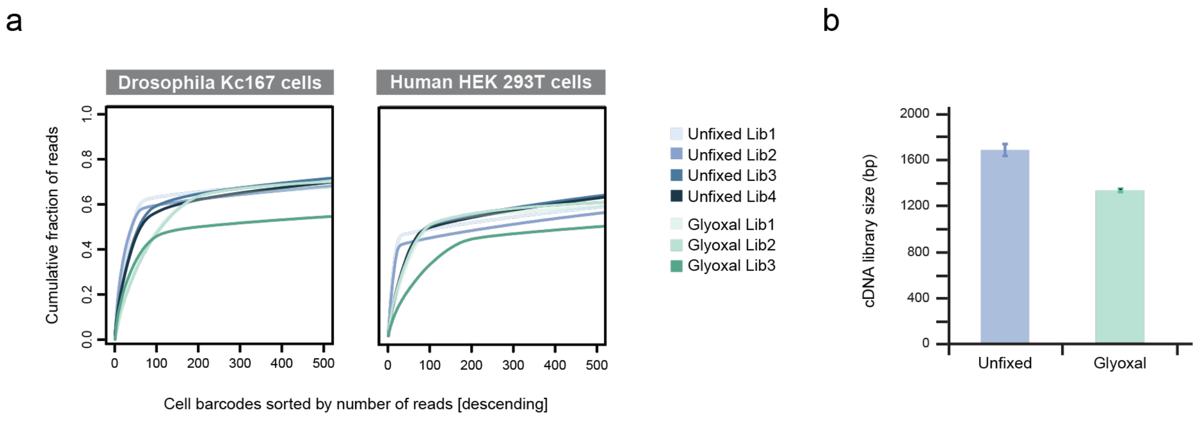


**Supplementary figure 1. Quality of single cell RNA-seq data. (a)** Kneeplots for *Drosophila* and human cell lines. The fraction of cumulative sequencing reads is plotted against cell barcodes sorted by descending number of sequencing reads. Inflection point of the data was used to estimate the number of "true" and "empty" cell barcodes. Four unfixed and three glyoxal-fixed technical replicates were analyzed for each cell line. **(b)** Average cDNA library size from unfixed and glyoxal-fixed mixed species samples depicted as bar graphs. Two technical and two biological replicates were measured per condition. Error bars indicate standard deviation between replicates.

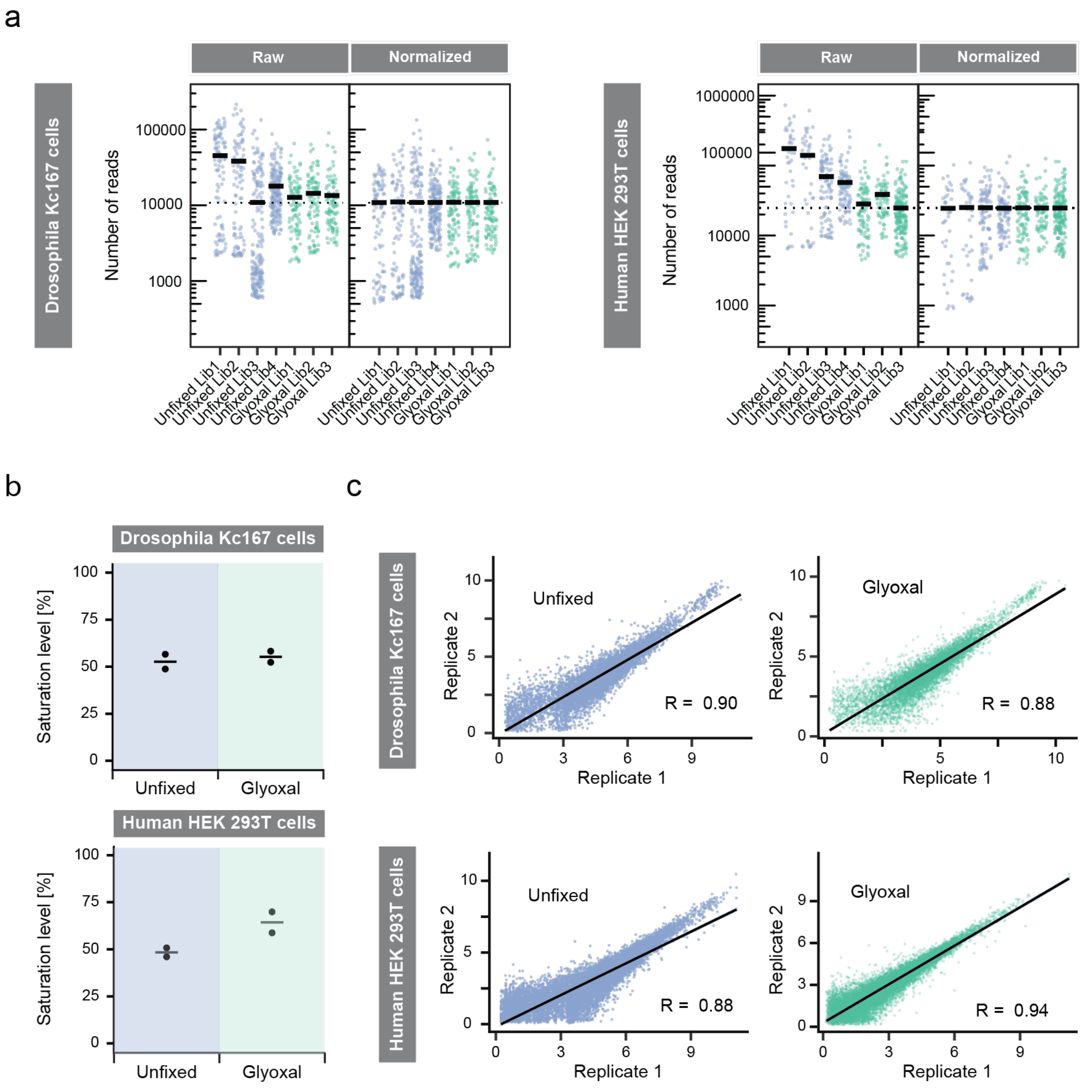


**Supplementary figure 2. Data normalization, library saturations and gene expression correlations between biological replicates. (a)** Sequencing depth before and after normalization from unfixed and glyoxal-fixed *Drosophila* and human cell lines. Horizontal line indicates average sequencing depth for each library. Data was normalized according to the average number of reads per cell among the different samples (dotted line). Four unfixed and three glyoxal-fixed technical replicates were analyzed for each cell line. **(b)** Assessment of saturation level of UMI numbers for unfixed and glyoxal-fixed single cell solutions obtained from *Drosophila* or human cell lines. **(c)** Average gene expression correlation of shared genes between biological replicates for unfixed and glyoxal-fixed *Drosophila* and human cell lines. Data is shown as normalized ATPM. R indicates Pearson correlation coefficient.

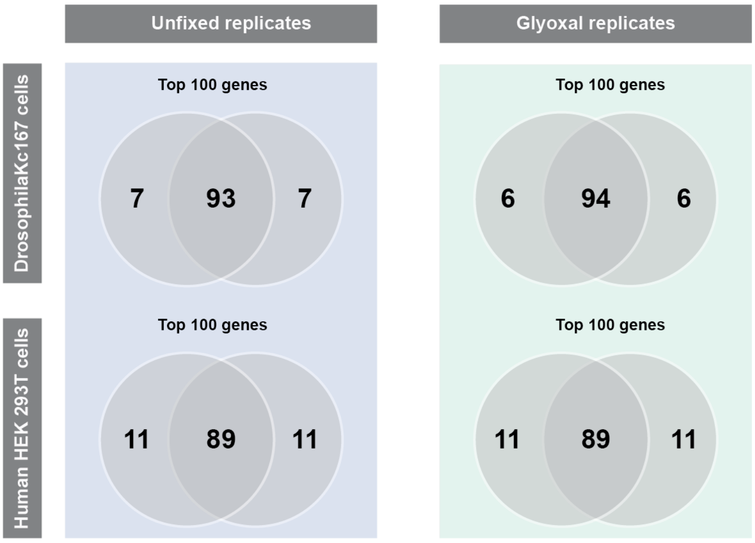


**Supplementary figure 3. Transcriptome comparison of unfixed and glyoxal-fixed *Drosophila* and human cells.** Overlap between top 100 highly expressed genes between technical replicates of unfixed and glyoxal-fixed *Drosophila* and human cell lines depicted in Venn diagrams.
